# Shoot-root interaction in control of camalexin exudation in Arabidopsis

**DOI:** 10.1101/2020.12.15.422875

**Authors:** Anna Koprivova, Vanessa Volz, Stanislav Kopriva

## Abstract

Plants exude secondary metabolites from the roots to shape the composition and function of their microbiome. Many of these compounds are known for their anti-microbial activity and are part of the plant immunity, such as the indole-derived phytoalexin camalexin. Here we studied the dynamics of camalexin synthesis and exudation upon induction of *Arabidopsis thaliana* with a plant growth promotion bacteria *Pseudomonas sp*. CH267 or a bacterial pathogen *Burkholderia glumae* PG1. We show that while the camalexin accumulation and exudation is more rapidly but transiently induced upon interaction with the growth promoting strain, the pathogen induces a higher and more stable camalexin levels. The concentration of camalexin in shoots, roots and exudates is well correlated, triggering a question on the origin of the exuded camalexin. By combination of experiments with cut shoots and roots and grafting of wild type plant with mutants in camalexin synthesis we showed that while camalexin can be produced and released by both organs, in intact plant the exuded camalexin originates in the shoots. We show that camalexin synthesis in response to *B. glumae* PG1 is dependent on cooperation of four CYP71 genes and a loss of function of any of them reduces camalexin synthesis. In conclusion, camalexin synthesis seems to be controlled on a whole plant level and coordinated between shoots and roots.

## INTRODUCTION

Plants are cohabiting their natural environments with plethora of microorganisms some beneficial or commensal, some harmful (Bulgarelli *et al*., 2013). Plants therefore evolved number of mechanisms that enable them to communicate with the microbiota, to attract the beneficial ones and defend themselves against the harmful ones. Number of these mechanisms are based on plant metabolites that can fulfil both of these functions (reviewed in (Jacoby *et al*., 2020; Sasse *et al*., 2018). Plants produce a number of secondary compounds that are directly involved in defense (Piasecka *et al*., 2015). Some of these compounds are synthesised only in response to the infection, and therefore they are classified as phytoalexins, whereas others are constitutive and activated upon tissue damage or pathogen triggered signalling; these are termed phytoanticipins (VanEtten *et al*., 1994). Chemically, the metabolites used by plants for defense belong to all major classes of secondary compounds, terpenes, phenolic compounds, and alkaloids (Zaynab *et al*., 2018).

One of the best characterised classes of phytoalexins are the sulfur containing indolic compounds, such as camalexin and brassinin in the crucifers (Pedras and Yaya, 2010). Camalexin, 3-thiazol-2′-yl-indole, accumulates upon infection with fungal pathogens, such as *Botrytis cinerea* or *Alternaria brassicicola* (Bednarek *et al*., 2005; Kliebenstein *et al*., 2005; Millet *et al*., 2010; Thomma *et al*., 1999). Variation in camalexin synthesis is associated with variation to susceptibility to *Botrytis* in Arabidopsis accessions (Rowe and Kliebenstein, 2008). Camalexin is synthesised from tryptophan, the first step in the pathway being the production of indole-3-acetaldoxime (IAOx), a common precursor for auxin, camalexin, and indole glucosinolate synthesis (Glawischnig *et al*., 2004). The first dedicated step in camalexin synthesis is the conversion of IAOx into indole-3-acetonitrile (IAN) by CYP71A12 and CYP71A13 (Glawischnig *et al*., 2004; Nafisi *et al*., 2007). IAN is conjugated by glutathione, which introduces the sulfur into the chemical structure and camalexin is ultimately synthesised by CYP71B15 (Geu-Flores *et al*., 2011; Schuhegger *et al*., 2006; Su *et al*., 2011; Zhou *et al*., 1999). The pathway may, however, be more complex, as two other P-450 enzymes, CYP71A27 and CYP71A28, were associated with camalexin accumulation in roots (Koprivova *et al*., 2019). The role of the individual isoforms particularly in roots is thus not very clear.

In the roots, camalexin was shown to have additional function to innate immunity, as a metabolite shaping the function of root associated microbiota (Koprivova *et al*., 2019). Using sulfatase activity in rhizosphere soil from Arabidopsis accessions as a measure for microbiome activity, genome wide association analysis showed that variation in CYP71A27 affects this microbial function. Loss of CYP71A27 resulted in lower sulfatase activity in soil, which could be complemented by camalexin. In addition, the *cyp71A27* mutant did not benefit from plant growth promoting (PGP) effects of several rhizospheric bacteria, which again could be complemented by addition of camalexin (Koprivova *et al*., 2019). Camalexin is exuded from the roots (Koprivova *et al*., 2019; Millet *et al*., 2010) and may represent an important player in the mechanisms by which plants control their microbiome (Jacoby *et al*., 2020). However, camalexin exudation seems to be in conflict with its definition as phytoalexin, as phytoalexins act in the site of their synthesis (VanEtten *et al*., 1994). Thus, it is important to discover more about the nature and control of camalexin exudation.

Here we show that camalexin exudation is triggered by both pathogenic and PGP bacteria and that camalexin accumulation in exudates, roots, and leaves is highly correlated. We also reveal that both leaves and roots are able to synthesise camalexin and used grafting to show that the camalexin exuded upon treatment of roots with *Burkholderia glumae* originates in the shoot.

## RESULTS

### Both PGP and pathogenic bacteria trigger camalexin synthesis and exudation

Previous work showed that camalexin can be exuded from plant roots incubated with PGP bacteria or the bacterial-derived peptide elicitor flagellin, which is the pre-requisite of camalexin function in shaping microbiome function (Koprivova *et al*., 2019; Millet *et al*., 2010). PGP bacteria and flagellin trigger also camalexin accumulation in roots, as does infection with root fungal pathogen *Verticillium longisporum* (Iven *et al*., 2012). To test, whether camalexin synthesis and exudation is triggered also by bacterial pathogens we incubated Arabidopsis growing in hydroculture with *Burkholderia glumae* PG1 (Gao *et al*., 2015) or a PGP bacterium *Pseudomonas sp*. CH267 (Haney *et al*., 2015; Koprivova *et al*., 2019). To obtain a better picture of a control of camalexin synthesis we analysed its accumulation in roots and shoots and in the exudates. Both bacteria triggered camalexin synthesis in all three compartments, whereas minimal camalexin levels were detected in roots and shoots of mock treated plants and no camalexin was exuded without the bacterial trigger (Figure 1). The two bacteria elicited camalexin synthesis and exudation in a different way, but similar in all three compartments. The PGP strain *Pseudomonas sp*. CH267 triggered a rapid response but the accumulation of camalexin peaked between 2 and 4 days and decreased afterwards, whereas the synthesis and exudation reached a maximum after 4 to 5 days and remained high upon treatment with *B. glumae* PG1. In the first days, the camalexin levels were higher upon treatment with *Pseudomonas sp*. CH267 but in later stages the pathogenic strain *B. glumae* PG1 triggered significantly higher levels of camalexin (Figure 1). Interestingly, even though the bacteria were in contact only with the roots, the camalexin concentrations were highly correlated between shoots and roots and also between both organs and the exudates. It is thus not possible to conclude whether the exuded camalexin is synthesised in the roots or the shoots.

**Figure 1.**
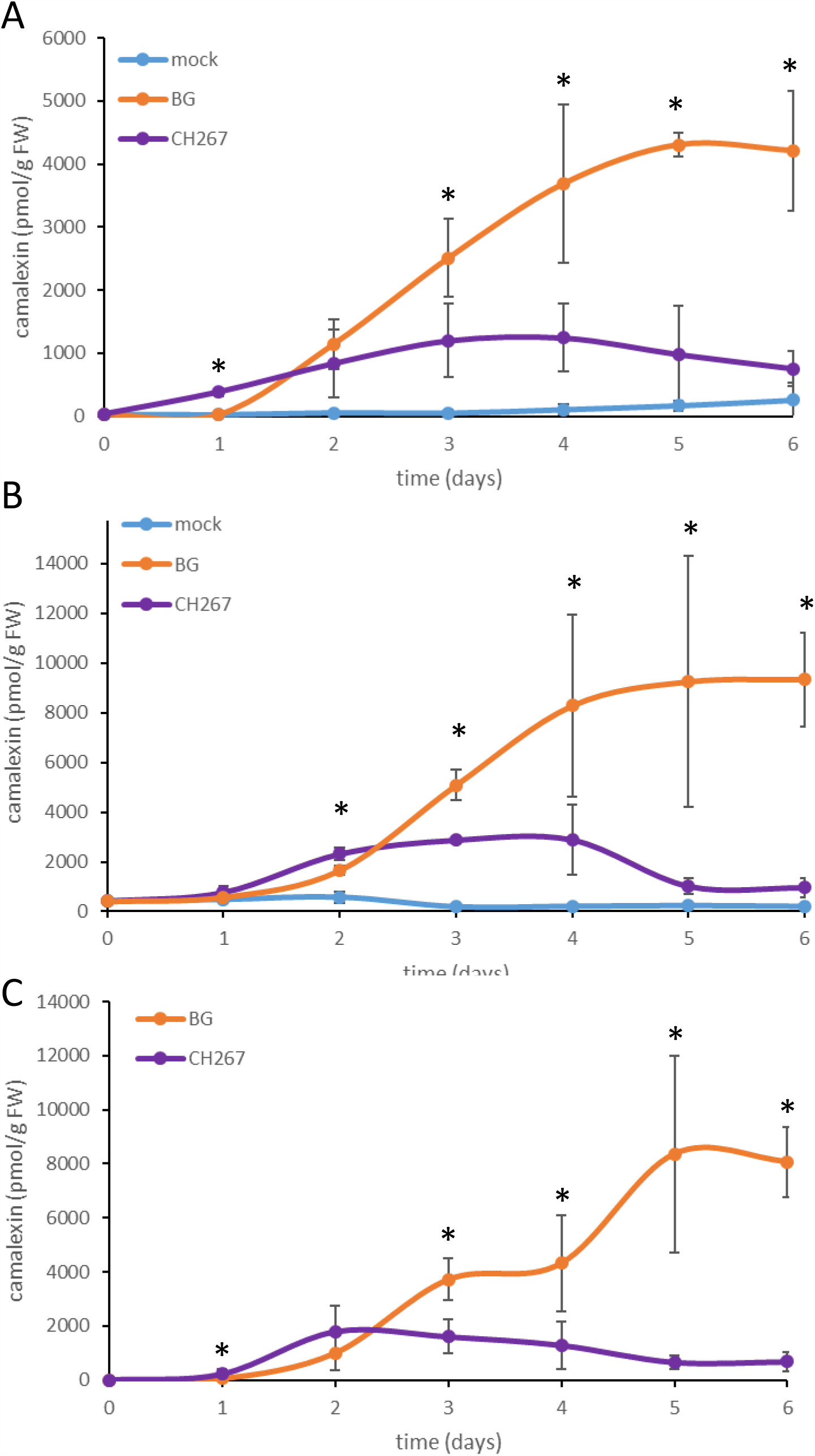
Camalexin accumulation upon inoculation with PGP or pathogen bacteria. Arabidopsis plants were grown on a nylon net in hydroculture for 10 days and inoculated in the solution with *Pseudomonas sp*. CH267, *B. glumae* PG1 (BG), or MgCl_2_ as mock. Camalexin was measured in leaves (**A**), roots (**B**), and exudates (**C**) sampled daily over 6 days. Data are presented as means ± S.D. from 4 biological replicates, each corresponding to at least 30 seedlings. Asterisks mark significant differences between the values of CH267 and PG1 treated plants (P<0.05, T-test).

### Contribution of different isoforms of CYP71A family to camalexin synthesis

We found previously that loss of two additional members of the CYP71A family of P-450 enzymes, CYP71A27 and CYP71A28, affected camalexin levels in roots (Koprivova *et al*., 2019). We were therefore interested in their contribution to total camalexin synthesis and obtained all possible double and triple mutants of the four isoforms CYP71A12, CYP71A13, CYP71A27, and CYP71A28. *CYP71A12* and *CYP71A13* as well as *CYP71A27* and *CYP71A28* are two pairs of neighbouring genes, and while a double mutant *cyp71a12 cyp71a13* (*cyp12/13*) has been produced by TALEN mutagenesis (Muller *et al*., 2015), double mutant of *CYP71A27* and *CYP71A28* is not available. These mutants were subjected to treatment with *B. glumae* PG1 for 3 days, leading to high synthesis of camalexin and allowing a good comparison of the individual genotypes. This analysis showed clearly that all four P-450 isoforms are important for camalexin synthesis (Figure 2A). Surprisingly, loss of CYP71A27 and CYP71A28 also led to a significant reduction of camalexin synthesis in the leaves, even if the corresponding genes are not expressed there. Camalexin levels in *cyp12/13* mutant is very low and in the range measured in mock treated plants, but still additional loss of either CYP71A27 or CYP71A28 lowers the camalexin accumulation further (Figure 2A). However, it needs to be seen, whether the effects of the mutations are due to loss of enzymatic activity or alteration of expression of other isoforms. Therefore, we determined the transcript levels of genes of camalexin synthesis pathway in all these mutants in roots. Inoculation with *B. glumae* PG1 led to increase of mRNA levels in roots of the genes for the enzymes of the canonical camalexin synthesis pathway CYP71A12, CYP71A13, and CYP7B15, as well as of *CYP71A27* (Figure 2B). As expected, the expression of the camalexin synthesis genes have been affected in the various mutants. The induction of *CYP71A12* by *B. glumae* PG1 was attenuated in the single mutants of other P-450 isoforms and in the double mutant *cyp13/27*, but surprisingly, increased in *cyp13/28* (Figure 2B). Also the induction of *CYP71B15* was less pronounced in the mutants. On the other hand, *CYP71A13* transcript levels were significantly elevated in genotypes with disrupted *CYP71A12* already without bacterial trigger. Although the *CYP71A28* mRNA was not detectable, disruption of this gene resulted in increased transcript levels of CYP71A27 both with and without inoculation (Figure 2B). Thus, the each of the four CYP71A isoforms seem to play some role in the camalexin network as loss of any of them affects at least one other member.

**Figure 2.**
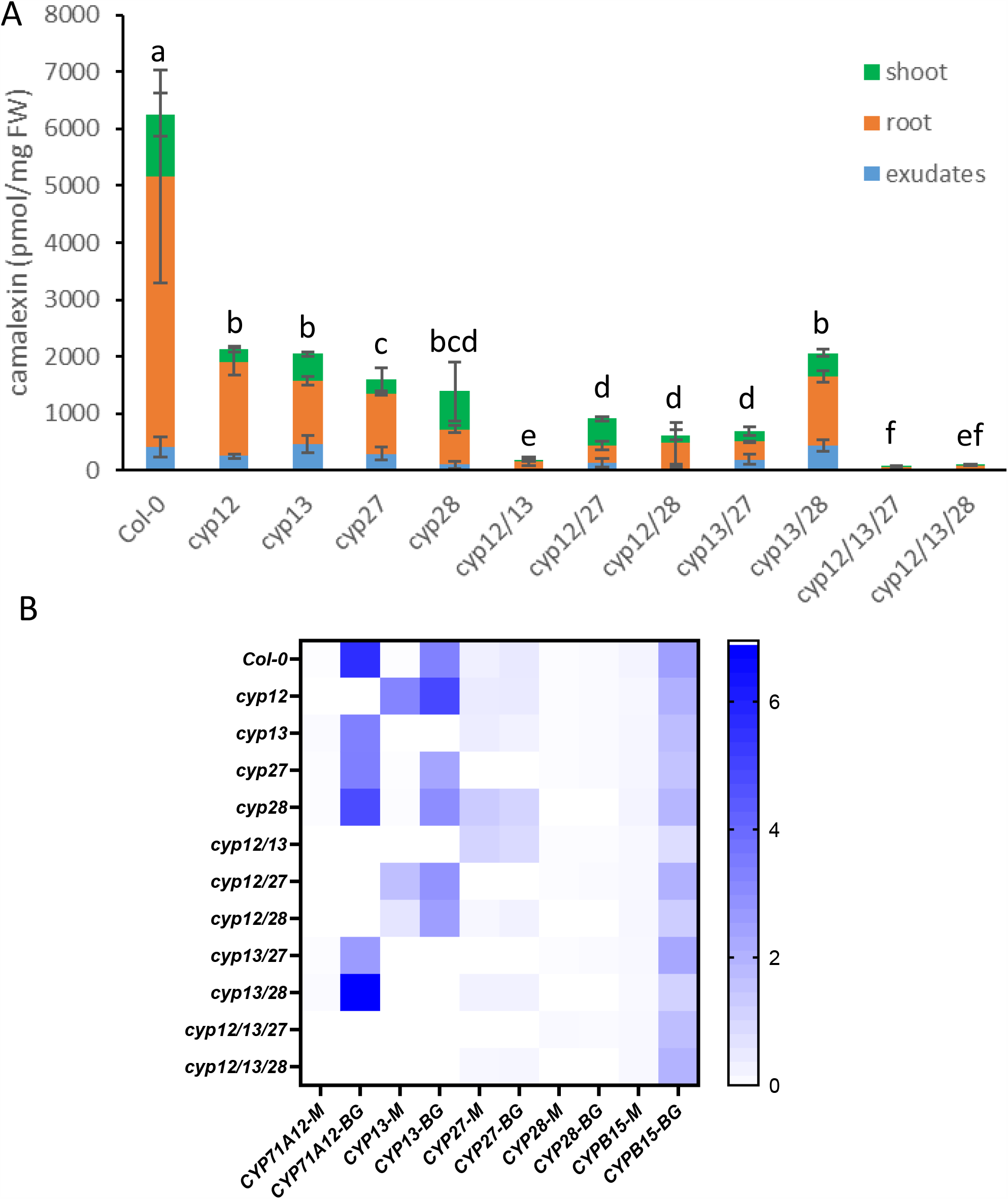
Characterisation of mutants in CYP71A genes involved in camalexin synthesis. The seedlings were grown on a nylon net in hydroculture for 10 days, inoculated in the solution with *B. glumae* PG1 or MgCl_2_ as mock and incubated for 3 days. **A** Camalexin was measured in leaves, roots, and exudates of *B. glumae* PG1 treated plants. Data are presented as means ± S.D. from 4 biological replicates, each corresponding to at least 30 seedlings. Different letters mark significant differences in total camalexin (shoots + roots + exudates) between the genotypes (P<0.05, ANOVA). **B** Transcript levels of the genes of camalexin synthesis were compared by RT-qPCR in roots of mock (M) and *B. glumae* PG1 (BG) treated plants. Data are shown as heatmap of relative expression.

### Dissection of tissue specificity of camalexin synthesis and exudation

The coordinated accumulation of camalexin in shoots, roots, and exudates after exposure of roots led to a question, whether the bacteria trigger camalexin synthesis also in the leaves. We, therefore, grew Arabidopsis plants on agarose plates, inoculated either the leaves or the root tips with the two bacterial strains and measured camalexin after 3 days incubation. Both inoculations triggered accumulation of camalexin in shoots and roots, but to a different extent depending on the bacterial strains. *Pseudomonas sp*. CH267 induced only a small camalexin accumulation, which did not differ neither in the two organs nor in the two types of inoculation and was only slightly higher than the levels found in sterile plants (Figure 3A). *B. glumae* PG1 triggered a similarly low camalexin synthesis when inoculated from root tip, but resulted in a large accumulation in leaves and to some extent also roots when inoculated onto leaves. Camalexin synthesis in Arabidopsis leaves thus react to *B. glumae* PG1 in the same way as to the fungal pathogens. The induction of camalexin synthesis in roots might be due to camalexin transport or to movement of the bacteria in the plant. We therefore used qPCR to determine bacterial titre in the plant material. No amplification was possible using primers for *Pseudomonas sp*. CH267, probably due to a low titre in our inoculations. Using primers for *B. glumae* PG1, however, bacteria were clearly detected in both roots and shoots, irrespective of the inoculated tissue, which reveals the mobility of this strain within the plant, both root-to-shoot and shoot-to-root directions (Figure 3B). Interestingly, while in plants inoculated from the root tip the amount of camalexin approximately correlates to the bacterial titre, in the plants inoculated from the leaves, the leaf camalexin concentration was almost 20-fold higher in leaf than in roots, despite a similar bacterial titre (Figure 3).

**Figure 3.**
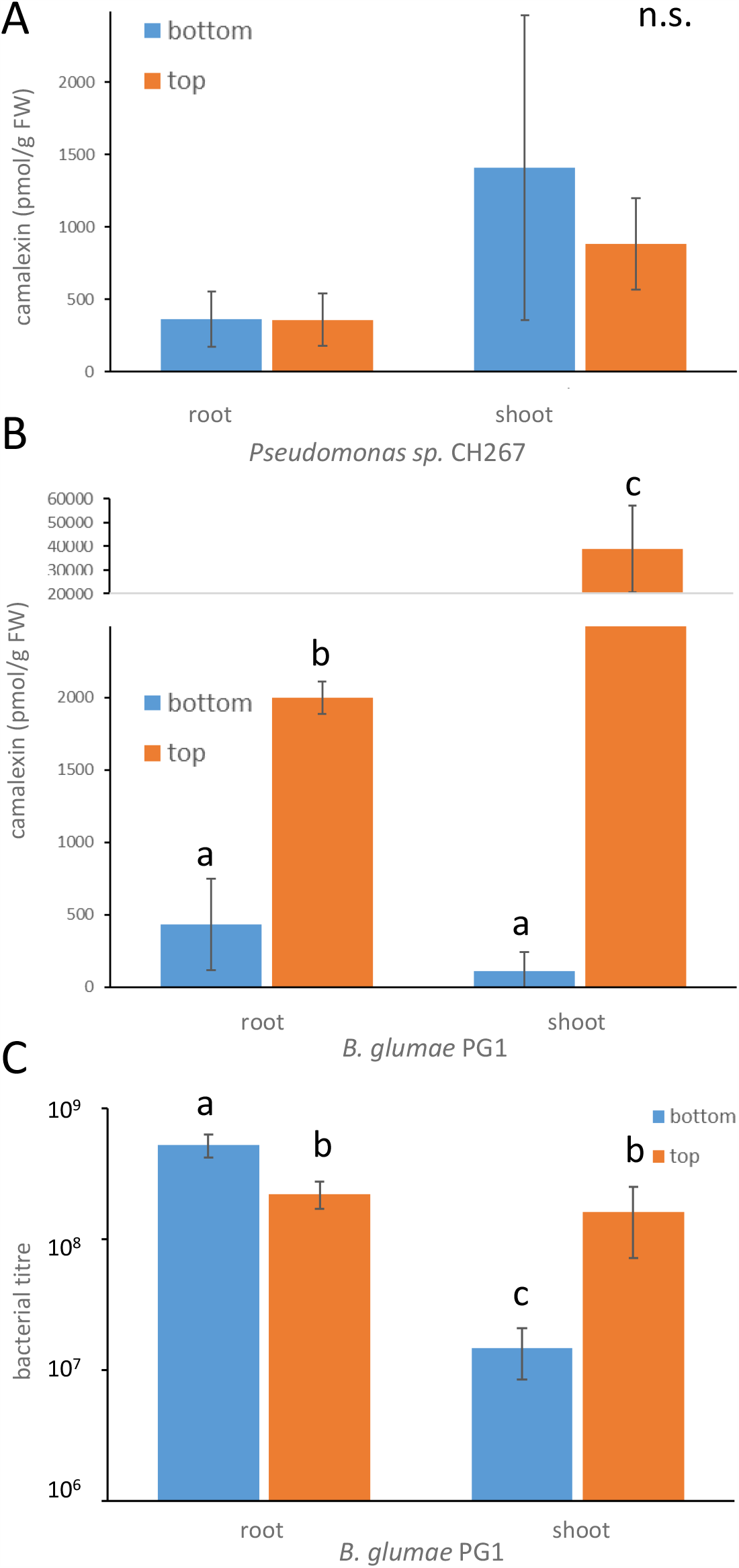
Tissue specificity of camalexin synthesis. Arabidopsis seedlings were grown for 14 days on an agar plate, inoculated with *Pseudomonas sp*. CH267 (**A)** or *B. glumae* PG1 (BG) (**B**) either on the leaves or on the root tips, and incubated for 3 days. Camalexin accumulation in leaves and roots was determined by HPLC. Data are presented as means ± S.D. from 3 biological replicates, each corresponding to 3 individual roots or shoots. **C** DNA was isolated from the roots and shoots and subjected to qPCR with primers against *B. glumae* PG1 and Arabidopsis TIP41 gene as control. Using previously established calibration between Ct values, OD and cfu, the qPCR data were expressed as cfu, presented as means ± S.D. from 4 biological replicates, each corresponding to 3 individual roots or shoots. Different letters mark values significantly different at P<0.05 (T-test).

We therefore asked, whether a communication between shoot and root affects camalexin synthesis in response to *B. glumae* PG1. We used the hydroponics system with plants growing on a nylon membrane, cut the shoots, placed them and the corresponding remaining roots separately in the wells of the 12 well plates, and inoculated with *B. glumae* PG1. Camalexin was then determined in the tissues and the exudates, as well as in shoot, root and exudates of intact plants analysed as controls (Figure 4). Both cut roots and shoots were able to exude camalexin to the solution, to levels higher than intact plants. Interestingly, whereas cut shoots accumulated more camalexin than shoots of intact plants, cut roots possessed only very low camalexin concentration compared to the intact controls. Total camalexin production in cut shoots was with 24±4 nmol g^-1^ FW higher than in intact plants (17±1 nmol g^-1^ FW) and cut roots (12±2 nmol g^-1^ FW). Thus, clearly, both roots and shoots are able to synthesise camalexin and its synthesis and exudation undergoes a control dependent on root-shoot communication.

**Figure 4.**
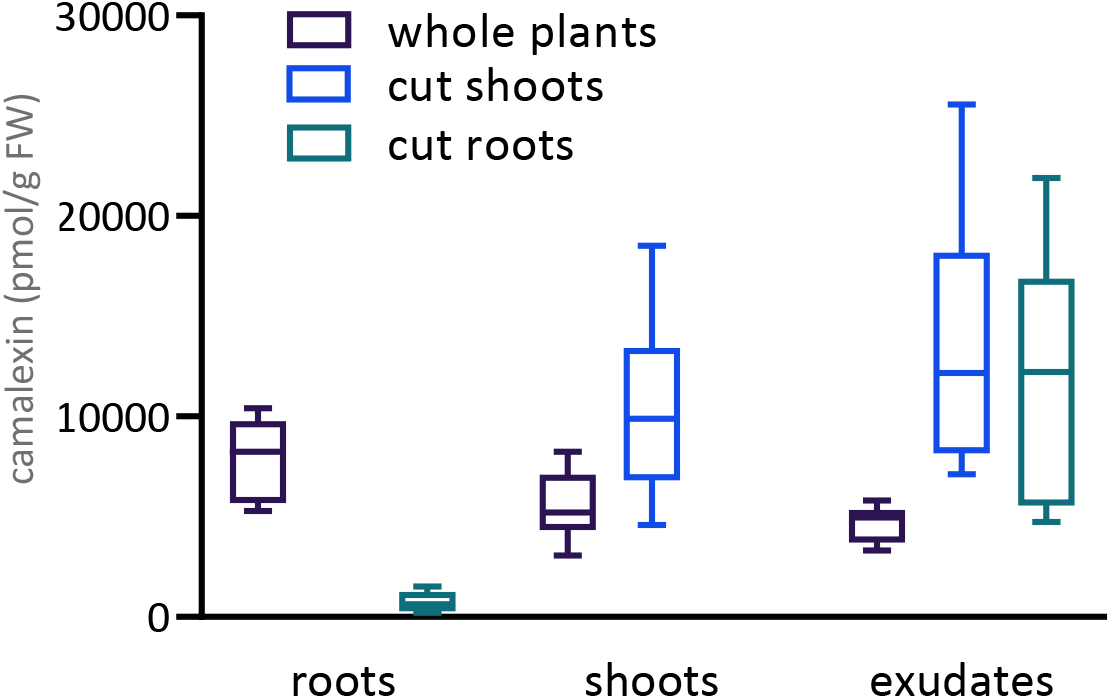
Camalexin in cut roots and shoots. Arabidopsis seedlings were grown on a nylon net in hydroculture for 10 days, the shoots were cut with scissors, the roots removed from the net and both placed separately to the solution. The shoots, roots, and intact plants were inoculated with *B. glumae* PG1 and further incubated for 3 days. Camalexin accumulation in shoots, roots, and exudates was determined by HPLC. Data are presented as box plots from at least 8 biological replicates, each corresponding to about 30 individual roots or shoots. The box extends from the 25^th^ to 75^th^ percentiles, the line is plotted at the median, the whiskers extend from minimum to maximum values.

While the experiments with cut roots and shoots were informative, they do not correspond to the *in vivo* situation. In order to determine where the camalexin exuded by root inoculation with *B. glumae* PG1 is synthesised, we performed grafting experiments with two mutants unable to synthesis camalexin, *pad3* and *cyp79b2 cyp79b3* (*b2/b3*) (Hull *et al*., 2000; Zhou *et al*., 1999). Inoculation of roots with *B. glumae* PG1 resulted in accumulation of camalexin in roots, shoots and exudates of Col-0 wild type (WT) homografts but not in homografts of the two mutants (Figure 5A). Even with a large variation due to analysis of individual seedlings, it can be clearly seen that in comparison with WT homografts, similarly high camalexin accumulation was found only in heterografted shoots originating from WT. Shoots of *b2/b3* mutant grafted on WT roots did not contain any camalexin higher that the background, whereas the shoots of *pad3* contained a low level of camalexin. Interestingly, WT roots grafted with *b2/b3* shoots contained same level of camalexin as roots of WT homografts, while WT roots grafted with *pad3* shoots did not contain any camalexin above the background. *pad3* roots grafted with WT shoots accumulated camalexin but *b2/b3* roots did not (Figure 5A). Importantly, camalexin was found only on exudates of grafted plants with WT shoots. Thus, the camalexin exuded upon inoculation of the roots by *B. glumae* PG1 must originate in the shoots.

**Figure 5.**
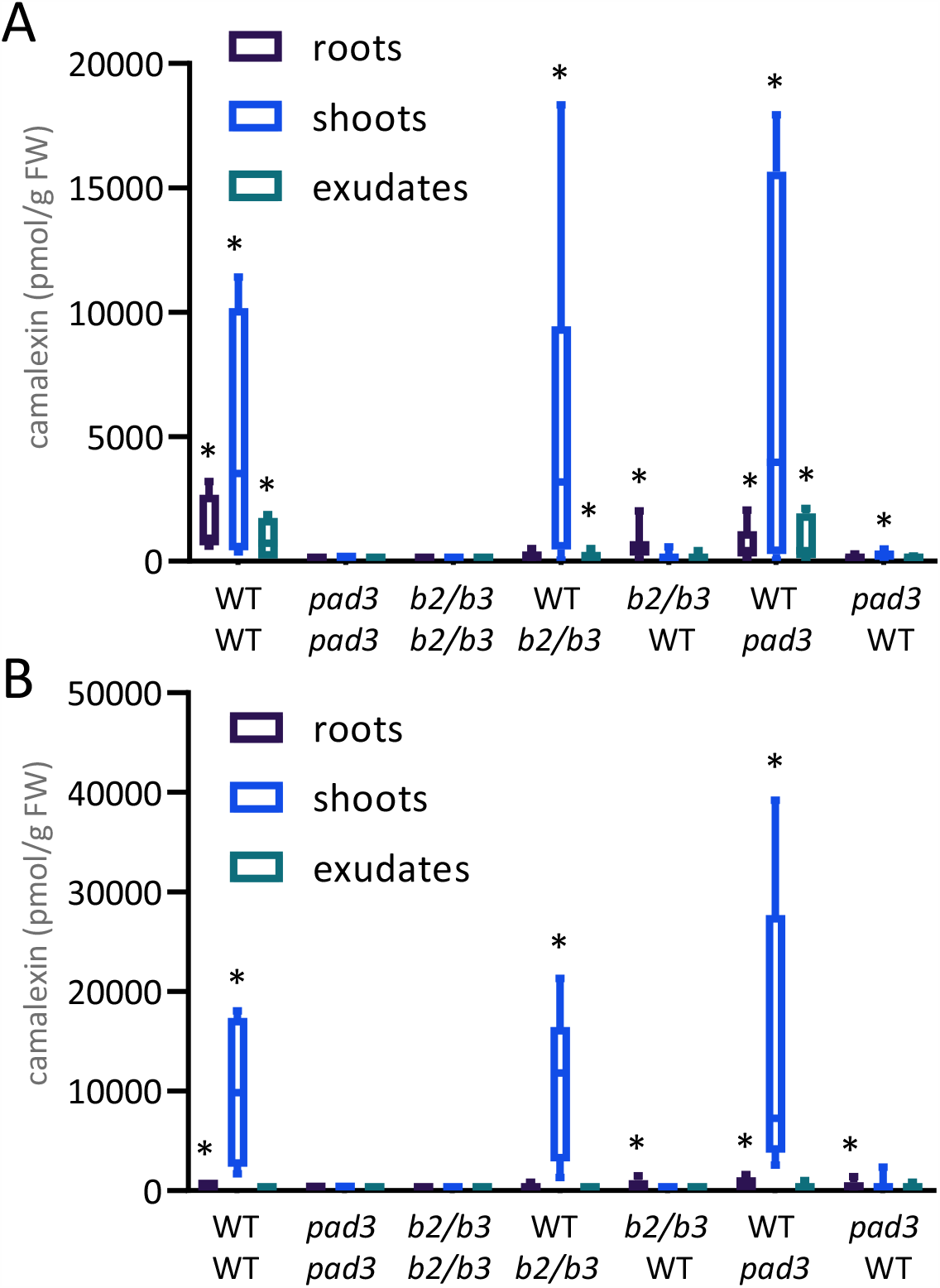
Analysis of camalexin in grafted plants. Homografts of Arabidopsis WT, *cyp79b2 cyp79b3* (*b2/b3*), and *pad3* and the heterografts of the WT with the mutants were grown for 18 after the grafting, transferred onto cut caps of Eppendorf tubes and placed with only the roots submerged into the hydroculture solution. The plants were then inoculated with B. glumae PG1 into the solution (**A**) or onto the leaves (**B**) and further incubated for 3 days. Camalexin accumulation in shoots, roots, and exudates was determined by HPLC. Data are presented as box plots from at least 8 individual grafts. The box extends from the 25^th^ to 75^th^ percentiles, the line is plotted at the median, the whiskers extend from minimum to maximum values.

When the grafted plants were inoculated with *B. glumae* PG1 on the leaves, camalexin was found mainly in shoots of WT homografts with much lower levels in roots and concentration not different to background in the exudates (Figure 5B). In the heterografts with *b2/b3*, only the tissues originating from WT accumulated camalexin, and none was exuded, whereas in grafts with *pad3* high level of camalexin was found in the shoots originating from WT, and low levels in both types of roots. Thus in whole plants camalexin synthesis seems to be tightly controlled and primarily occurring in the shoots.

## DISCUSSION

Camalexin is a relatively new addition to the list of plant metabolites that shape the root associated microbiome (Jacoby *et al*., 2020; Koprivova *et al*., 2019). Similar to other such compounds, e.g. coumarins or benzoxazinoids (de Bruijn *et al*., 2018; Stringlis *et al*., 2018), camalexin has first been characterised for its antimicrobial properties (Rogers *et al*., 1996). However, in the rhizosphere these compounds affect the microbiota in a way that they support plant fitness and performance. For example, coumarins exuded from plants were shown to affect the communities to improve plant iron nutrition (Harbort *et al*., 2020). Camalexin was shown to affect the microbial sulfatase activity in rhizosphere soil that mineralises organic sulfur and so the bacteria help plants to access this sulfur pool (Kertesz and Mirleau, 2004; Koprivova *et al*., 2019). Thus, camalexin has to be exuded to fulfil this function, however, unlike the coumarins or benzoxazinoids, camalexin has not been found in roots exudates unless elicited (Millet *et al*., 2010; Monchgesang *et al*., 2016; Neal *et al*., 2012). In addition, while the other metabolites characterised so far change the taxonomic assembly of the microbial community, this still needs to be tested for camalexin.

Since camalexin has always been a prime example of a phytoalexin, i.e. acting locally at the site of pathogen attack, it is not obvious how camalexin exudation from the roots is regulated. Previous work focused on camalexin synthesis in the leaves as a reaction to leaf pathogens (Glazebrook and Ausubel, 1994; Kliebenstein *et al*., 2005; Thomma *et al*., 1999). It was shown previously that the exudation could be elicited by flagellin or by PGP bacteria (Koprivova *et al*., 2019; Millet *et al*., 2010), but to relatively low levels. It was therefore important that high camalexin exudation can be triggered by the pathogenic bacteria *B. glumae* PG1 as a robust high exudation is needed to dissect the regulation. Interestingly, the dynamics of camalexin synthesis and exudation responds differentially to pathogenic and commensual or beneficial bacteria. While the PGP strain seemed to trigger the camalexin synthesis quicker, the response was only transient. The pathogenic bacterial strain seemed to be slower in initiation of the synthesis, but this was stronger and remained active for longer and did not diminish. The levels of camalexin in roots of *B. glumae* PG1 treated Arabidopsis plants in our system were similar to those in roots treated with *V. longisporum* (Iven et al., 2012). It seems therefore, that while the camalexin synthesis is initiated by both types of microorganisms, only upon interaction with pathogens the synthesis and exudation is sustained long-term. Similarly, when the plants were grown on agar plates and inoculated with from the leaves, *B. glumae* PG1 induced much higher camalexin levels than *Pseudomonas sp*. CH267 (Figure 3). *B. glumae* PG1 can thus be used as a tool to study the control of camalexin synthesis and exudation.

The first question addressed using *B. glumae* PG1 was the contribution of the “old” (CYP71A12 and A13) and “new” (A27 and A28) CYP71A isoforms to camalexin synthesis and exudation. All mutants clearly showed reduced total accumulation of camalexin, when the concentrations in shoots, roots, and exudates were summed (Figure 2A). This was particularly true for the concentration in the roots, which contributed most to the total camalexin; not surprisingly, as the pathogen was inoculated by the roots. However, with exception of *cyp71a28* all other mutants showed also reduced accumulation in the shoots. This result is in contrast to previous experiments with the “old” *cyp71a12* and *cyp71a13*, as in the former, upon abiotic elicitation with UV light or AgNO_3_camalexin in leaves was not affected and upon treatment with spores of fungal pathogen *Plectosphaerella cucumerina BMM* even increased (Muller *et al*., 2015; Pastorczyk *et al*., 2020). On the other hand it agrees with the measurements of camalexin in roots of soil grown plants, where all four single mutants showed lower concentrations (Koprivova *et al*., 2019). The data also clearly demonstrate the very high contribution of CYP71A12 and CYP71A13 to camalexin synthesis in all compartments, despite previous conclusion that CYP71A12 is responsible for root synthesis and exudation upon elicitation with flagellin (Millet *et al*., 2010). However, the data also show that even in the absence of these two enzymes, some camalexin is produced and this production is dependent on CYP71A27. Interestingly, the loss of the individual genes affects also transcript levels of the other members of the biosynthesis network (Figure 2B). With one notable exception, an induction of *CYP71A13* in the *cyp71a12* background, the level of induction of the genes by *B. glumae* PG1 was attenuated in the mutants. Since the enzymes of camalexin synthesis form a metabolon (Mucha *et al*., 2019) this might be a mechanism to prevent accumulation of proteins that cannot be part of this structure.

The analysis of camalexin in the 3 compartments of different mutants showed that its concentrations correlate well between all three of them. The genes for camalexin synthesis are also expressed in both shoots and roots. Thus, the camalexin in the exudates might originate in the roots as well as in the leaves.

To find out which organ is responsible for the synthesis of camalexin found in exudates we designed two experiments. In a simple approach we cut shoots and roots and incubated them separately with the bacterial pathogen (Figure 4). This experiment revealed that both shoots and roots are autonomous in camalexin synthesis. Shoots alone even produced more camalexin than the whole plants, that can be explained by the direct leakage of the synthesised camalexin and also by a more rapid contact of the shoots with the bacteria. The inoculation of leaves with *B. glumae* PG1 (Figure 3) resulted in a much higher camalexin accumulation in leaves compared to the hydroculture setup (Figures 1 and 2), which is consistent with the high camalexin in cut shoots. The high camalexin production in the cut roots, on the other hand, was unexpected, since in previous experiments with inoculation from the leaves the same bacterial titre that triggered accumulation of ca. 40 nmol mg^-1^ FW in leaves induced only 1 nmol mg^-1^ FW in the roots. This means that a coordination between shoots and roots is necessary to prevent camalexin overproduction. There are number of examples how roots and shoots communicate in defense, from the resistance against leaf pathogens induced by the rhizobacterium *Pseudomonas fluorescens* SS101, which also involves camalexin (van de Mortel *et al*., 2012), to the coordination in jasmonate signalling for resistance to nematodes (Wang *et al*., 2019). Camalexin synthesis can be affected by auxin and by miRNA393, both known long-distance signals (Robert-Seilaniantz *et al*., 2011). The nature of the signal controlling root camalexin synthesis in response to *B. glumae* PG1, however, still needs to be determined.

The existence of such coordination was clearly demonstrated in the second approach, using grafting with mutants that do not synthesise camalexin, *pad3* and *cyp79b2 cyp79b3*. Any camalexin found in the grafted tissues originating from the mutants must be transported from WT and thus evidence for a long-distance transport. Indeed, the grafting experiment showed unequivocally that camalexin exuded from the roots originates in the shoots. Camalexin was found in amounts over background only in exudates from heterografts with WT shoots, but not roots. Interestingly, when the plants were inoculated onto the leaves, camalexin synthesis in the leaves was induced to the same degree, but none of this camalexin was exuded. Thus the plants seem to recognise where the infection originates and steer the camalexin synthesis there. This process requires a sophisticated coordination between the roots and the shoots, and cannot rely solely on the actual perception of the bacteria, as seen also in Figure 3, where the same bacterial titres triggered different camalexin levels in shoots and roots. It remains to be seen whether camalexin exudation in response to PGP bacteria undergoes the same whole plant regulation or whether the camalexin is produced locally, in the root, as might be indicated by the lower production and different dynamics.

In conclusion, here we show that inoculation of Arabidopsis root with a bacterial pathogen *B. glumae* PG1 triggers camalexin synthesis in shoots and roots and its exudation. The camalexin can be produced and released by both organs, but in intact plant the exuded camalexin originates in the shoots. We show that the camalexin synthesis genes are tightly regulated and loss of function of any of them affects total camalexin synthesis. Finally, we conclude that camalexin synthesis is controlled by a whole plant regulation with a need for shoot root communication.

## MATERIALS AND METHODS

### Plant material and growth conditions

*Arabidopsis thaliana L*. accession Col-0 was used as wild type alongside mutants in camalexin synthesis *cyp71a12* (GABI_127H03), *cyp71a13* (SALK_105136), *pad3* (SALK_026585), *cyp71a27* (SALK_053817) and *cyp71a28* (SALK_ 064792). The double mutants *cyp71a12 cyp71a13* and *cyp79b2 cyp79b3* were obtained from H. Frerigmann and T. Gigolashvili, University of Cologne, respectively.

For camalexin and expression analyses plants were surface sterilized with chlorine gas. Seeds were suspended in 0.1 % agarose, distributed onto square 1 cm x 1 cm sterile nylon membranes (about 30 seeds per sample) and placed in 12 well plates on top of 1 ml of ½ Murashige Skoog (MS) medium with 0.5 % sucrose. After stratification for 2 days in dark and cold the plates were transferred to 22°C and kept in dark for 3 days to promote etiolation, which greatly simplifies the separation of shoots from membranes. Afterwards the plates were incubated at long day conditons (16 h light/ 8 h dark), 120 µE m^-2^ S^-1^, and at 22°C for further 7 days. The medium was then replaced with ½ MS without sucrose and the plants incubated for 24 hours before inoculation with the bacteria or mock and incubated further in the same conditions for 3 days, unless specified otherwise.

For the experiments with cut shoots and roots immediately before inoculation the shoots were cut with scissors and roots were freed from the membrane and placed directly into the nutrient solution.

### Bacterial strains and conditions for cocultivation experiments

For co cultivation experiments 2 bacterial strains were used, *Pseudomonas* sp. CH267 (Haney *et al*., 2015), obtained from J. R. Dinneny, Stanford University and *B. glumae* PG1 (Gao *et al*., 2015), obtained from K.-E. Jä ger, Heinrich Heine Universitä t Düsseldorf, Germany. The bacteria were kept as glycerol stocks and plated freshly before experiment on LB plates supplemented with appropriate antibiotics.

For inoculation, overnight bacterial cultures were washed two times with sterile 10 mM MgCl_2_ and final OD_600_ was measured. *Pseudomonas* sp. CH267 was diluted stepwise to OD_600_ = 0.0001, and *B. glumae* PG1 to OD_600_ = 0.0005 in 10 mM MgCl_2_. Eight µl of these suspensions were used for inoculation into each well. Eight µl of 10 mM MgCl_2_ was used as mock treatment. Samples for DNA, RNA and camalexin (shoots, roots and exudates) were harvested after 3 days of inoculation, except the time course experiments.

Alternatively, plants were grown on square Petri dishes with ½ MS with sucrose for 18 days and inoculated with 8 µl of suspensions of *Pseudomonas* sp. CH267 (OD_600_ = 0.0001) or *B. glumae* PG1 (OD_600_ = 0.0005) onto leaves or the bottom 2 mm of root tips. After 30 min drying the plates were returned to growth cabinet and grown for 3 days at long days.

### Camalexin measurements

Camalexin was extracted from 5-30 mg of plant material as described in (Koprivova *et al*., 2019). For extraction of camalexin from exudates the media were centrifuged at maximum speed for 20 min at 18°C and purified using 1 ml solid phase extraction tubes (Discovery-DSC18) according to manufacturer’s instructions. Samples were eluted with 90% (V/V) acetonitrile and 0.1% (V/V) formic acid, dried in a speed vac and dissolved in 50 µl of DMSO. 20 µl was injected into HPLC and analysed as described above. For the quantification external standards were used ranging from 1 pg to 1 ng per µl.

### Grafting experiments

For grafting, the selected genotypes were first grown on vertical Petri dishes containing ½ MS with sucrose and 0.8% agarose for 5-6 days at short day conditions (10 h light / 14 h dark) and 22°C. Grafting was performed under binocular microscope in sterile conditions and the grafts were transferred onto fresh plates for further 18 days under the same short day light conditions. Graft unions were examined under the binocular to identify adventitious root formation. Healthy grafts were carefully transferred into 12 well plates with 1 ml of ½ MS medium placing the shoots onto sterile cut cups from 0.5 ml Eppendorf tubes in order to prevent them from direct contact with the liquid. The plants were inoculated with 8 µl of *B. glumae* PG1 suspension at OD_600_ = 0.0005 into the solution or onto the leaves and further incubated for 3 days. Camalexin analysis in shoots, roots, and exudates of individual plants was performed as described above.

### Expression analysis

To determine transcript levels total RNA was isolated by standard phenol/chlorophorm extraction and LiCl precipitation. First strand cDNA synthesis was performed using QuantiTect Reverse transcription Kit (Quiagen) from 800 ng of total RNA. Quantitative real time RT-PCR (qPCR) was performed using gene-specific primers (Supplemental Table 1) and the fluorescent dye SYBR Green (Promega). All quantifications were normalized to the TIP41 (AT4G34270) gene. The RT-PCR reactions were performed in duplicate for each of the 4 independent samples.

### Determination of bacterial titre

For the estimation of bacterial titre using qPCR the method from (Ross and Somssich, 2016) was adapted. Genomic DNA was extracted using buffer containing 0.025 M EDTA, 0.2 M Tris pH 8.0, 0.25 M NaCl and 0.5% SDS. After 10 min incubation at 65°C and subsequent centrifugation, supernatant was precipitated with equal volume of isopropanol, washed with 70% ethanol and resuspended in 100 µl of sterile water. For the qPCR 13 ng of corresponding DNA samples were used with *Arabidopsis* (At primer AT4G26410) and *B. glumae* PG1 specific primer (Burk1 for NR042931). The qPCR conditions were the same as for expression analysis. The qPCR reactions were performed in duplicate for each of the 4 independent samples. To relate the qPCR results to the bacterial titre, first serial dilutions of bacterial suspensions of different OD_600_ have been plated on LB plates and the colonies were counted manually to link OD_600_ and cfu. Subsequently, 10 µl of five 10-fold dilutions of bacterial suspensions with initial OD_600_ = 1.8 were added to 30 mg of Arabidopsis leaves and the DNA extracted and analysed as above. Using calibration curves plotting ΔCt (Ct_Bg_ - Ct_At_) and the cfu against the log_10_OD_600_ the bacterial titre can be estimated from the ΔCt values.

## ACKNOWLEDGEMENTS

Research in SK’s lab is funded by the Deutsche Forschungsgemeinschaft (DFG) under Germany’s Excellence Strategy – EXC 2048/1 – project 390686111 and within the SPP 2125 DECRyPT.

## References

Bednarek P, Schneider B, Svatos A, Oldham NJ, Hahlbrock K. 2005. Structural complexity, differential response to infection, and tissue specificity of indolic and phenylpropanoid secondary metabolism in Arabidopsis roots. Plant Physiology 138, 1058–1070.

Bulgarelli D, Schlaeppi K, Spaepen S, Ver Loren van Themaat E, Schulze-Lefert P. 2013. Structure and functions of the bacterial microbiota of plants. Annual Review of Plant Biology, Vol 62 64, 807–838.

de Bruijn WJC, Gruppen H, Vincken JP. 2018. Structure and biosynthesis of benzoxazinoids: Plant defence metabolites with potential as antimicrobial scaffolds. Phytochemistry 155, 233–243.

Gao R, Krysciak D, Petersen K, Utpatel C, Knapp A, Schmeisser C, Daniel R, Voget S, Jaeger KE, Streit WR. 2015. Genome-wide RNA sequencing analysis of quorum sensing-controlled regulons in the plant-associated Burkholderia glumae PG1 strain. Appl Environ Microbiol 81, 7993–8007.

Geu-Flores F, Moldrup ME, Bottcher C, Olsen CE, Scheel D, Halkier BA. 2011. Cytosolic gamma-glutamyl peptidases process glutathione conjugates in the biosynthesis of glucosinolates and camalexin in Arabidopsis. Plant Cell 23, 2456–2469.

Glawischnig E, Hansen BG, Olsen CE, Halkier BA. 2004. Camalexin is synthesized from indole-3-acetaldoxime, a key branching point between primary and secondary metabolism in Arabidopsis. Proc Natl Acad Sci U S A 101, 8245–8250.

Glazebrook J, Ausubel FM. 1994. Isolation of phytoalexin-deficient mutants of Arabidopsis thaliana and characterization of their interactions with bacterial pathogens. Proc Natl Acad Sci U S A 91, 8955–8959.

Haney CH, Samuel BS, Bush J, Ausubel FM. 2015. Associations with rhizosphere bacteria can confer an adaptive advantage to plants. Nat Plants 1, 15051.

Harbort CJ, Hashimoto M, Inoue H, Niu Y, Guan R, Rombola AD, Kopriva S, Voges M, Sattely ES, Garrido-Oter R, Schulze-Lefert P. 2020. Root-Secreted Coumarins and the Microbiota Interact to Improve Iron Nutrition in Arabidopsis. Cell Host Microbe 28, 825–837 e826.

Hull AK, Vij R, Celenza JL. 2000. Arabidopsis cytochrome P450s that catalyze the first step of tryptophan-dependent indole-3-acetic acid biosynthesis. Proc Natl Acad Sci U S A 97, 2379–2384.

Iven T, Konig S, Singh S, Braus-Stromeyer SA, Bischoff M, Tietze LF, Braus GH, Lipka V, Feussner I, Droge-Laser W. 2012. Transcriptional activation and production of tryptophan-derived secondary metabolites in arabidopsis roots contributes to the defense against the fungal vascular pathogen Verticillium longisporum. Molecular Plant 5, 1389–1402.

Jacoby RP, Koprivova A, Kopriva S. 2020. Pinpointing secondary metabolites that shape the composition and function of the plant microbiome. Journal of Experimental Botany.

Kertesz MA, Mirleau P. 2004. The role of soil microbes in plant sulphur nutrition. Journal of Experimental Botany 55, 1939–1945.

Kliebenstein DJ, Rowe HC, Denby KJ. 2005. Secondary metabolites influence Arabidopsis/Botrytis interactions: variation in host production and pathogen sensitivity. Plant Journal 44, 25–36.

Koprivova A, Schuck S, Jacoby RP, Klinkhammer I, Welter B, Leson L, Martyn A, Nauen J, Grabenhorst N, Mandelkow JF, Zuccaro A, Zeier J, Kopriva S. 2019. Root-specific camalexin biosynthesis controls the plant growth-promoting effects of multiple bacterial strains. Proc Natl Acad Sci U S A 116, 15735–15744.

Millet YA, Danna CH, Clay NK, Songnuan W, Simon MD, Werck-Reichhart D, Ausubel FM. 2010. Innate immune responses activated in Arabidopsis roots by microbe-associated molecular patterns. Plant Cell 22, 973–990.

Monchgesang S, Strehmel N, Schmidt S, Westphal L, Taruttis F, Muller E, Herklotz S, Neumann S, Scheel D. 2016. Natural variation of root exudates in Arabidopsis thaliana-linking metabolomic and genomic data. Sci Rep 6, 29033.

Mucha S, Heinzlmeir S, Kriechbaumer V, Strickland B, Kirchhelle C, Choudhary M, Kowalski N, Eichmann R, Huckelhoven R, Grill E, Kuster B, Glawischnig E. 2019. The Formation of a Camalexin Biosynthetic Metabolon. Plant Cell 31, 2697–2710.

Muller TM, Bottcher C, Morbitzer R, Gotz CC, Lehmann J, Lahaye T, Glawischnig E. 2015. TRANSCRIPTION ACTIVATOR-LIKE EFFECTOR NUCLEASE-Mediated Generation and Metabolic Analysis of Camalexin-Deficient cyp71a12 cyp71a13 Double Knockout Lines. Plant Physiology 168, 849–858.

Nafisi M, Goregaoker S, Botanga CJ, Glawischnig E, Olsen CE, Halkier BA, Glazebrook J. 2007. Arabidopsis cytochrome P450 monooxygenase 71A13 catalyzes the conversion of indole-3-acetaldoxime in camalexin synthesis. Plant Cell 19, 2039–2052.

Neal AL, Ahmad S, Gordon-Weeks R, Ton J. 2012. Benzoxazinoids in root exudates of maize attract Pseudomonas putida to the rhizosphere. PLoS One 7, e35498.

Pastorczyk M, Kosaka A, Pislewska-Bednarek M, Lopez G, Frerigmann H, Kulak K, Glawischnig E, Molina A, Takano Y, Bednarek P. 2020. The role of CYP71A12 monooxygenase in pathogen-triggered tryptophan metabolism and Arabidopsis immunity. New Phytologist 225, 400–412.

Pedras MS, Yaya EE. 2010. Phytoalexins from Brassicaceae: news from the front. Phytochemistry 71, 1191–1197.

Piasecka A, Jedrzejczak-Rey N, Bednarek P. 2015. Secondary metabolites in plant innate immunity: conserved function of divergent chemicals. New Phytologist 206, 948–964.

Robert-Seilaniantz A, MacLean D, Jikumaru Y, Hill L, Yamaguchi S, Kamiya Y, Jones JD. 2011. The microRNA miR393 re-directs secondary metabolite biosynthesis away from camalexin and towards glucosinolates. Plant Journal 67, 218–231.

Rogers EE, Glazebrook J, Ausubel FM. 1996. Mode of action of the Arabidopsis thaliana phytoalexin camalexin and its role in Arabidopsis-pathogen interactions. Mol Plant Microbe Interact 9, 748–757.

Ross A, Somssich IE. 2016. A DNA-based real-time PCR assay for robust growth quantification of the bacterial pathogen Pseudomonas syringae on Arabidopsis thaliana. Plant Methods 12, 48.

Rowe HC, Kliebenstein DJ. 2008. Complex genetics control natural variation in Arabidopsis thaliana resistance to Botrytis cinerea. Genetics 180, 2237–2250.

Sasse J, Martinoia E, Northen T. 2018. Feed Your Friends: Do Plant Exudates Shape the Root Microbiome? Trends in Plant Science 23, 25–41.

Schuhegger R, Nafisi M, Mansourova M, Petersen BL, Olsen CE, Svatos A, Halkier BA, Glawischnig E. 2006. CYP71B15 (PAD3) catalyzes the final step in camalexin biosynthesis. Plant Physiology 141, 1248–1254.

Stringlis IA, Yu K, Feussner K, de Jonge R, Van Bentum S, Van Verk MC, Berendsen RL, Bakker P, Feussner I, Pieterse CMJ. 2018. MYB72-dependent coumarin exudation shapes root microbiome assembly to promote plant health. Proc Natl Acad Sci U S A 115, E5213–E5222.

Su T, Xu J, Li Y, Lei L, Zhao L, Yang H, Feng J, Liu G, Ren D. 2011. Glutathione-indole-3-acetonitrile is required for camalexin biosynthesis in Arabidopsis thaliana. Plant Cell 23, 364–380.

Thomma BP, Nelissen I, Eggermont K, Broekaert WF. 1999. Deficiency in phytoalexin production causes enhanced susceptibility of Arabidopsis thaliana to the fungus Alternaria brassicicola. Plant Journal 19, 163–171.

van de Mortel JE, de Vos RC, Dekkers E, Pineda A, Guillod L, Bouwmeester K, van Loon JJ, Dicke M, Raaijmakers JM. 2012. Metabolic and transcriptomic changes induced in Arabidopsis by the rhizobacterium Pseudomonas fluorescens SS101. Plant Physiology 160, 2173–2188.

VanEtten HD, Mansfield JW, Bailey JA, Farmer EE. 1994. Two Classes of Plant Antibiotics: Phytoalexins versus “Phytoanticipins”. Plant Cell 6, 1191–1192.

Wang G, Hu C, Zhou J, Liu Y, Cai J, Pan C, Wang Y, Wu X, Shi K, Xia X, Zhou Y, Foyer CH, Yu J. 2019. Systemic Root-Shoot Signaling Drives Jasmonate-Based Root Defense against Nematodes. Curr Biol 29, 3430–3438 e3434.

Zaynab M, Fatima M, Abbas S, Sharif Y, Umair M, Zafar MH, Bahadar K. 2018. Role of secondary metabolites in plant defense against pathogens. Microb Pathog 124, 198–202.

Zhou N, Tootle TL, Glazebrook J. 1999. Arabidopsis PAD3, a gene required for camalexin biosynthesis, encodes a putative cytochrome P450 monooxygenase. Plant Cell 11, 2419–2428.

