## Supplemental Table 1 for "Shoot-root interaction in control of camalexin exudation in Arabidopsis"

Supplemental Table 1. Primers used in the study

| | | Gene ID | Forward | Reverse |
|---|---|---|---|---|
| 1 | *TIP41* | **AT4G34270** | gaactggctgacaatggagtg | atcaactctcagccaaaatcg |
| 2 | *CYP71A12* | **AT2G30750** | tgtggtgtttggtccctatg | ttgttcgtgagcagattgaga |
| 3 | *CYP71A13* | **AT2G30770** | gatgttgtgtttgctccctatg | ttgttggtgagcagattgaga |
| 4 | *CYP71A27* | **AT4G20240** | ccctacggagaagattggaa | ccagcttctctgtcattactttga |
| 5 | *CYP71A28* | **AT4G20235** | ttctcctcctacggcgaata | gaggagatggacagtgcataaa |
| 6 | *CYP71B15* | **AT3G26830** | caccactgatcatctcaaagga | cggtcattccccatagtgtt |
| 7 | *At* | **AT4G26410** | gagctgaagtggcttccatgac | ggtccgacatacccatgatcc |
| 8 | *Burk1* | **NR042931** | ggaactgcatttgtgactgg | ctccccacgctttcgtgc |
